# TAD Evolutionary and functional characterization reveals diversity in mammalian TAD boundary properties and function

**DOI:** 10.1101/2023.03.07.531534

**Authors:** Mariam Okhovat, Jake VanCampen, Ana C. Lima, Kimberly A. Nevonen, Cora E. Layman, Samantha Ward, Jarod Herrera, Alexandra M. Stendahl, Ran Yang, Lana Harshman, Weiyu Li, Rory R. Sheng, Yafei Mao, Lev Fedorov, Blaise Ndjamen, Katinka A. Vigh-Conrad, Ian R. Matthews, Sarah A. Easow, Dylan K. Chan, Taha A. Jan, Evan E. Eichler, Sandra Rugonyi, Donald F. Conrad, Nadav Ahituv, Lucia Carbone

**Author notes:** These authors equally contributed to the project (Mariam Okhovat, Jake VanCampen, Ana C. Lima).

## Abstract

Topological associating domains (TADs) are self-interacting genomic units crucial for shaping gene regulation patterns. Despite their importance, the extent of their evolutionary conservation and its functional implications remain largely unknown. In this study, we generate Hi-C and ChIP-seq data and compare TAD organization across four primate and four rodent species, and characterize the genetic and epigenetic properties of TAD boundaries in correspondence to their evolutionary conservation. We find that only 14% of all human TAD boundaries are shared among all eight species (ultraconserved), while 15% are human-specific. Ultraconserved TAD boundaries have stronger insulation strength, CTCF binding, and enrichment of older retrotransposons, compared to species-specific boundaries. CRISPR-Cas9 knockouts of two ultraconserved boundaries in mouse models leads to tissue-specific gene expression changes and morphological phenotypes. Deletion of a human-specific boundary near the autism-related *AUTS2* gene results in upregulation of this gene in neurons. Overall, our study provides pertinent TAD boundary evolutionary conservation annotations, and showcase the functional importance of TAD evolution.

## INTRODUCTION

The three-dimensional (3D) organization of the genome plays a fundamental role in orchestrating chromatin interactions that regulate gene expression and shape phenotypes^1,2^. Chromatin conformation capture assays, namely Hi-C, have shed light on 3D genome organization and revealed that chromosomes are compartmentalized into kilo-to mega-base scale segments termed topologically associated domains (TADs). TADs often contain gene(s) that can interact with regulatory elements located in the same domain, while interactions across domains are prevented by flanking elements that are often bound by specific proteins including CTCF, cohesin complexes and RNA-polymerase II^3^. These insulating elements are commonly known as TAD boundaries and considering their significant role in defining functional interaction domains, they are thought to be critical for maintaining proper genome function.

Disruption of TAD boundaries has been associated with ectopic gene interactions, gene mis-regulation and aberrant phenotypes such as cancer^4^, limb malformation^5^ and neurodevelopmental disease^6^. As a result, the function and position of TAD boundaries in the genome are subject to some degree of evolutionary constraint and conservation. When TADs were first described^7–9^, a direct comparison between human and mouse TAD organization revealed that a high portion of TAD boundaries was shared between the two species (i.e., 53.8% of human boundaries were found to be boundaries in mouse)^7^. This finding was corroborated by several subsequent lines of direct and indirect evidence. For example, direct comparisons of the *Hox*^10^ and *Six*^11^ loci detected similar TAD structures across several species. Further convincing —albeit indirect— evidence of TAD conservation came from studies providing evidence for selection against structural variations (SVs) that disrupt genome topology; In the human population, evolutionary SVs were shown to be depleted at TAD boundaries, while this pattern was absent in patients with developmental delay^12^. In addition, in the highly rearranged gibbon genome^13^ breakpoints of evolutionary chromosomal rearrangements between human and gibbon were found to be enriched at TAD boundaries, resulting in preservation of TADs even when synteny is lost^14^. Findings from these and other studies have reinforced the notion that TAD boundaries remain unchanged during evolution, due to deleterious consequences of TAD disruption^15^. However, most of the evidence supporting TAD conservation is indirect and/or only limited to a few species or loci. Moreover, a few recent studies present evidence refuting high conservation of TAD boundaries across species, even those as closely related as human and chimpanzee^16^. Hence, there is growing uncertainty around the level of evolutionary TAD conservation^17^ and a need for further reassessment of TAD conservation via comprehensive direct cross-species comparisons.

Gene regulation plays a key role in species evolution^18^. Therefore, comparative investigation of TAD boundaries and the gene regulatory domains they form can shed new light on mechanisms contributing to adaptations, speciation, and evolution. Regions of high TAD conservation represent crucial loci whose disruption may lead to reduced fitness^19^, while regions with evolutionarily diverged TAD organization may be associated with gene interaction/expression changes that underlie evolutionary novelties across species. Indeed, differences in 3D genomic interactions in human-chimpanzee induced pluripotent stem cells (iPSC) have been associated with inter-specific differences in gene expression^16^. Furthermore, 3D chromatin structures identified in the fetal human cortex, but not in rhesus and mouse, may have contributed to human-specific features of brain development^20^, highlighting the power of using a comparative approach in assessing TAD evolution and inferring TAD boundary functionality.

Here, in order to assess the level of evolutionary TAD conservation in mammals, we identified TAD boundaries across four primate and four rodent species and performed direct comparisons to group them based on their evolutionary conservation. Characterization of the genomic and epigenetic profiles of TAD boundary across evolutionary groups, as well as generation of *in vivo* and *in vitro* deletions allowed us to investigate function of these boundaries and their relevance to development, disease and evolution.

## RESULTS

### Cross-species comparison of TAD boundaries reveals variation in evolutionary conservation

In order to identify TADs and their boundaries across species, we selected eight species from the primate and rodent orders, and generated genome-wide chromatin conformation (Hi-C) maps (**Fig. 1a**). The selected species all had high-quality reference genomes and spanned different evolutionary distances within their order. Furthermore, for both orders we included species that have experienced several evolutionary genomic rearrangements in a relatively short evolutionary time. For primates, in addition to human (representing great apes) and rhesus macaque (representing Old World Monkeys), we included two species of gibbons, as these small apes have significantly rearranged genomes between genera despite having diverged only ~5 million years ago^13^. The two selected gibbon species have very different karyotypes and chromosome numbers (*Nomascus leucogenys*, 2n=52 and *Hylobates moloch*, 2n=44), referred to as Nomascus and Hylobates hereon. From rodents, we selected mouse and rat, as well as two relatively closely related *Mus* species with divergent karyotypes (*M. Caroli*, 2n=40 and *M. pahari*, 2n=48), referred to as Caroli and Pahari from hereon^21^. We used public Hi-C data for human lymphoblastoid cell line (LCL) (**Supplementary Table 1**), but generated new high-resolution Hi-C data for all other species from one male and one female and merged the two replicates after confirming that they were highly correlated (*r*=0.66-0.75; (**Supplementary Table 1**, **Extended Data Fig. 1a**). Moreover, we generated CTCF and histone (H3K4me1, H3K27ac, H3K4me3 and H3K27me3) ChIP-seq data from the same samples (**Supplementary Table 2**). For rhesus and all rodent species, we used liver tissue. However, due to the challenge of sampling from endangered gibbons, we used LCL from these species.

**Fig 1.**
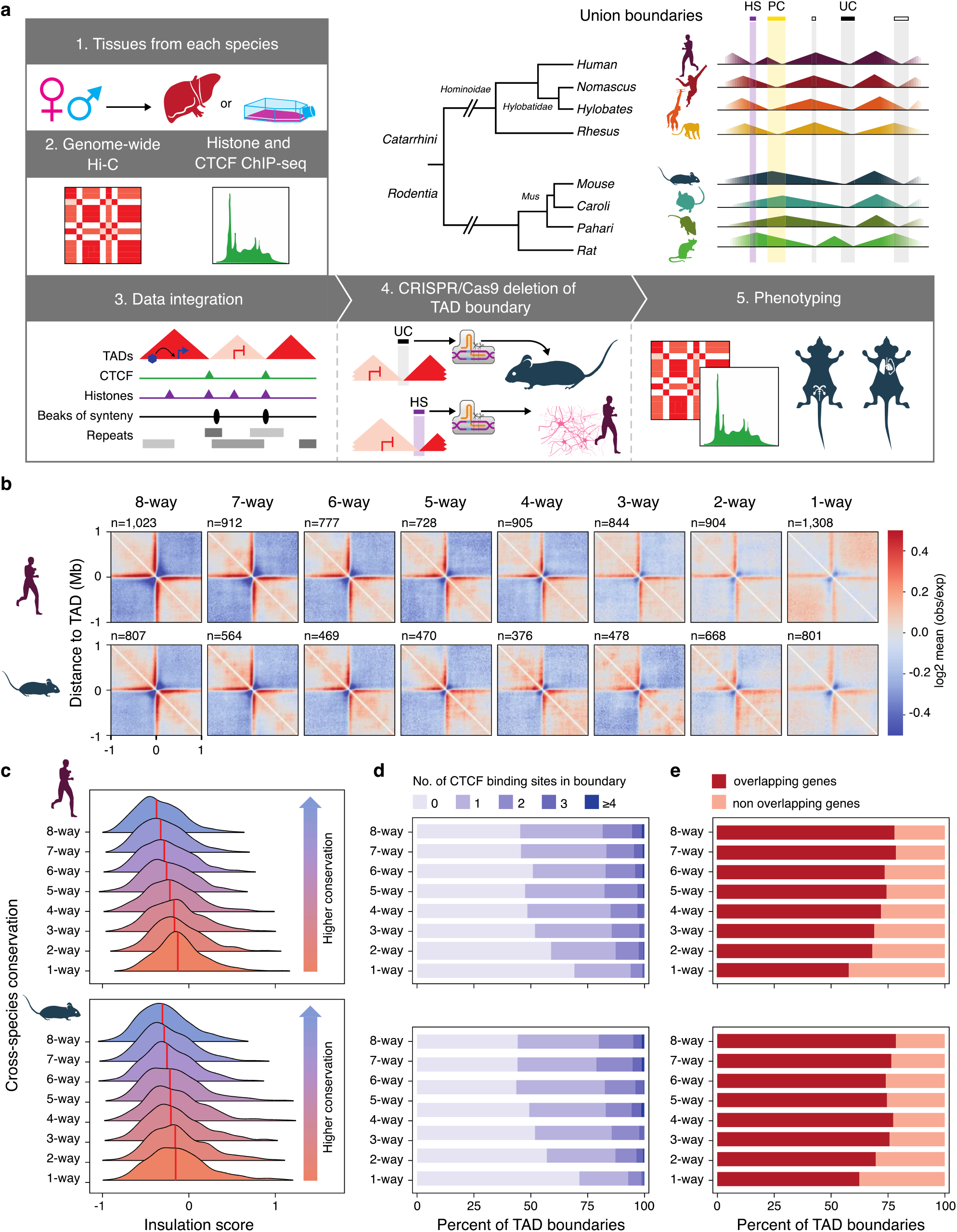
Direct multi-species comparisons shed light on evolutionary conservation of TAD boundaries. **a**, Flowchart of the study design is shown, along with a scheme of the union boundary approach and our evolutionary criteria for annotating human-specific (HS), primate-conserved (PC) and ultraconserved (UC) TAD boundaries. **b**, Heatmaps show frequency of genomic interactions (log2 mean observed/expected) across TAD boundaries increase as the TAD boundary conservation decrease from 8-to 1-way in both human and mouse genomes. **c**, Kernel density plots (median marked with red line) show lower insulation scores (i.e., stronger TAD separation) as boundary conservation increases. D, Percent TAD boundaries overlapping CTCF binding sites and genes are positively associated with the number of species sharing the boundary.

Using our Hi-C data, we identified an average of 7,177 ± 481 (mean ± stdev) TADs across the genomes of all species, with mouse (N=6,542) and rhesus (N=7,845) having the lowest and highest numbers of predicted TADs, respectively (**Supplementary Table 1**). Across all species and tissues, TAD boundaries exhibited local enrichment of CTCF and two histone marks associated with active transcription, H3K27ac and H3K4me3 (**Extended Data Fig. 1b**). Over half of TAD boundaries (62.2% ± 5.5; mean ± stdev) overlapped genes in all species, with the highest and lowest percent overlap present in human (70.7%) and Pahari (56.4%), respectively. On average, less than half (40 ± 5 %; mean ± stdev) of TAD boundaries in all species contained at least one CTCF binding site, with mouse having the highest (48%) and Nomascus having the lowest (36%) percentages (**Extended Data Fig. 2**).

### Genetic and epigenetic properties of boundaries vary depending on their evolutionary conservation

TADs have likely emerged at different evolutionary times and been exposed to different degrees of selective pressures. Hence, the evolutionary conservation and cross-species stability of TAD boundaries in a genome may vary greatly. In order to assess evolutionary conservation of TAD boundary placement across species, we used a “union boundary” strategy (**Supplementary Notes**) in which boundaries from each species were lifted over to the human genome coordinates (hg38, **Supplementary Table 3**) and merged within 10Kb of each other to create a union boundary reference map (**Supplementary Table 4**). Boundaries from all species were then compared against each union boundary, and those present in the human and mouse reference genomes were grouped based on the number of species sharing them (*e.g*., 1-way, 2-way, 3-way…., 8-way), so that the genetic and epigenetic features of these groups could be compared within their respective genome. Comparison of the overall Hi-C interaction matrices at boundaries in each boundary group in human and mouse genomes, revealed that interaction frequencies were overall lower across boundary groups that had higher cross-species conservation (**Fig. 1b**). To better quantify this, we also estimated the insulation score^22^ of TAD boundaries, which is calculated by summing contacts along the Hi-C matrix using sliding windows (**Methods**). We observed that as the level of conservation increased, the insulation scores decreased, indicating that boundaries shared by more species lead to higher separation between neighboring TADs (**Fig. 1c**). In agreement with this finding, boundaries shared by more species overlapped CTCF binding sites and genes more frequently than less conserved ones (**Fig. 1d-e**). Overall, these observations indicate that regardless of tissue and species, functional and structural properties of TAD boundaries differ depending on their cross-species conservation.

We then used the phylogenetic relationship between species to classify human and mouse TAD boundaries based on their evolutionary conservation and estimated age, into the following groups: 1) ultraconserved boundaries (examples in **Extended Data Fig. 3**), which are shared across all species in this study and date back to at least the common ancestor of primates and rodents (~80 mya); 2) primate-conserved or rodent-conserved, which are only shared within each order and are likely at least ~25 mya and ~15 mya old respectively; and 3) species-specific, which are only present in the mouse (mouse-specific) or the human (human-specific) genome and represent younger boundaries that have recently evolved in each genome (~7 mya for human and ~3 mya for mouse). Furthermore, to exclude human boundaries that are shared with other great apes, we used public Hi-C data from chimpanzee and gorilla LCL^23^. Based on our classifications, 13.6% (N=1,023) of the boundaries in the human LCL genome were ultraconserved, 6% (N=491) were primate-conserved, and 15% (N=1,130) were human-specific. A roughly similar distribution was found in the mouse liver genome, where 19.1% (N=1,023) of the TAD boundaries were ultraconserved, 2.1% (N=115) were rodent-conserved, and 14.2% (N=761) were mouse-specific (**Supplementary Table 4**).

We then made use of the species-specific epigenetic and genetic data to investigate properties of TAD boundaries as a function of their conservation in mouse and human genomes. We found that less conserved boundary groups in both human and mouse genomes showed significantly lower insulation scores and less overlap with genes, compared to older and more conserved boundary groups (Wilcoxon rank sum test, p<0.001; **Fig. 2a-b**). As differences in interaction patterns and insulation strength of TAD boundary groups could, at least partially, be attributed to their different frequency of overlap with genes, we also showed that insulation scores of both ultraconserved and species-specific TAD boundaries in mouse and human were significantly more extreme than randomly selected boundaries with the same frequency of overlap with genes (one-tail permutation p-value <0.05) (**Supplementary Notes**). The epigenetic landscape was also variable across boundary groups, with species-specific TAD boundaries in both human and mouse genomes showing higher enrichment of chromatin states associated with active transcription start sites, bivalent chromatin, and CTCF signal (**Fig. 2c**). Consistently, ultraconserved boundaries contained significantly more CTCF binding sites compared to order-conserved and species-specific boundaries in both the human and mouse genomes (Wilcoxon rank sum test, p<0.001) (**Fig. 2d**). Hence, regardless of species and tissue, boundaries stratified based on evolutionary conservation differ in their genetic and epigenetic properties, with ultraconserved boundaries displaying stronger insulation and higher dependence on CTCF binding compared to species-specific boundaries.

**Fig 2.**
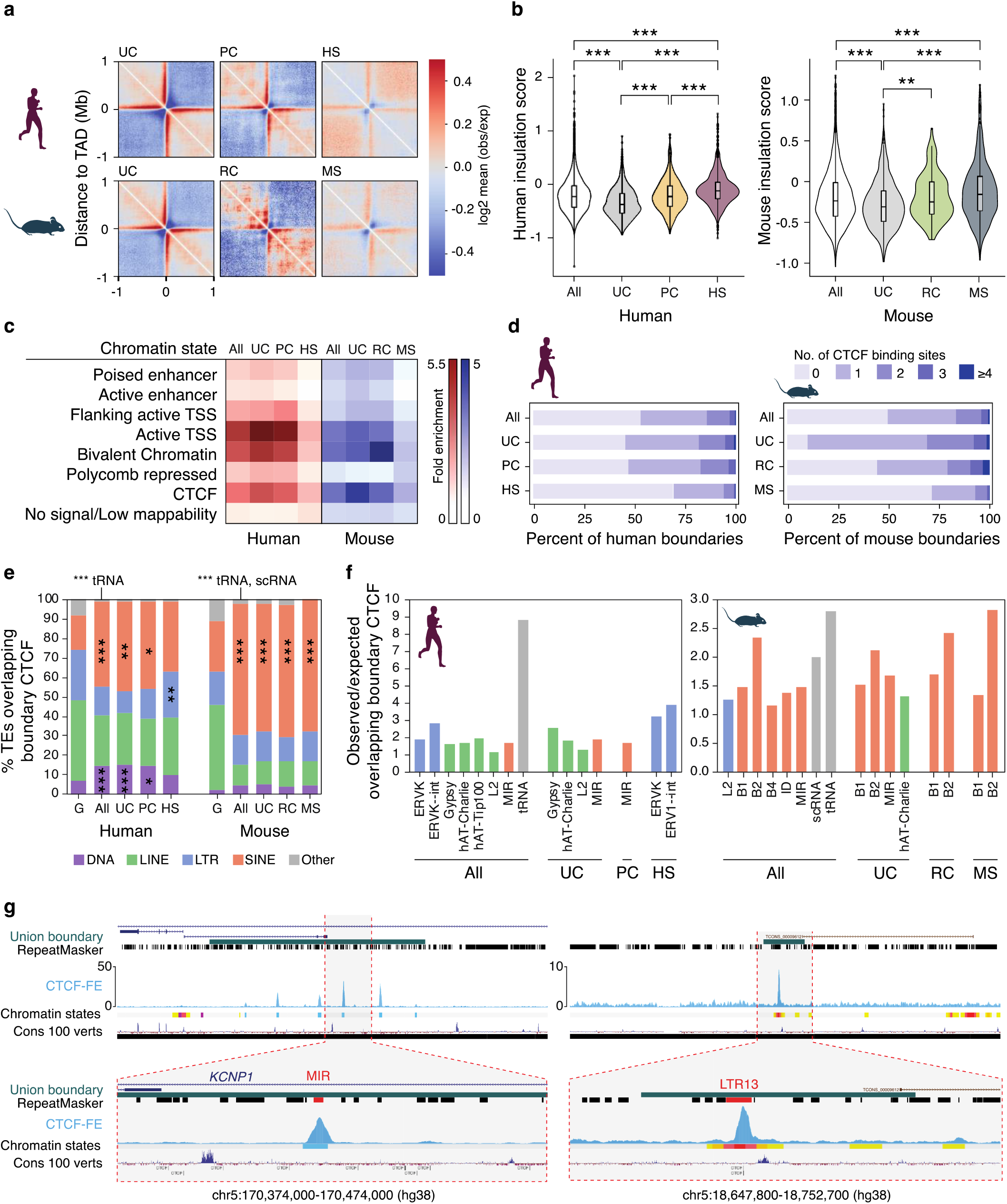
Genetic and epigenetic features of TAD boundaries vary as a function of their evolutionary conservation. **a**, Heatmaps of Hi-C interaction show differences in genomic interactions across evolutionary TAD boundary groups. **b**, Violin plots show insulation score differences across evolutionary TAD boundary groups in human (left) and mouse (right). P-values are only reported when <0.05. **c**, Fold-enrichment of chromatin states at evolutionary TAD boundary groups in human (red) and mouse genome (blue). **d**, Percent of TAD boundaries harboring CTCF binding sites across evolutionary groups. **e**, Percent CTCF binding sites in TAD boundaries that overlap TE classes in human (left) and mouse (right). **f**, Observed/Expected values for TE families significantly over-represented at TAD boundary CTCF binding sites. **g**, UCSC genome browser screenshots of a MIR-derived CTCF binding site inside an ultraconserved boundary (left), and an ERV-derived CTCF binding site within human-specific TAD boundary (right). CTCF ChIP-seq fold-enrichment (CTCF-FE) track appears in blue. G= genome-wide, All= All TAD boundaries, US= ultraconserved, PC= primate-conserved, HS= human-specific RC= rodent-conserved MS= mouse-specific; *** p<0.0005, ** p <0.005, * p<0.05.

### Transposable elements contribute to the evolution of conserved and species-specific TAD boundaries

Transposable elements (TEs) can introduce CTCF binding sites upon their insertion in host genomes^24^ and as such, their co-option may have contributed to the evolution of TAD boundaries^25,26^. Analysis of CTCF binding sites located inside TAD boundaries in each species, showed that over half of them overlap TEs across species (ranging from 53-77%, depending on species; **Extended data Fig. 2**). In the human genome, we found significant enrichment (binomial test; p<0.005) of SINE and DNA repeat classes at CTCF binding sites inside ultraconserved and primate-conserved boundaries (**Fig. 2e**).

Among the enriched SINE repeats, the most note-worthy family was MIR, a family of ancient t-RNA-derived and highly conserved TEs^27,28^. MIR elements have been previously shown to provide insulation in the human genome in a CTCF-independent manner^29^, but we found 14% of ultraconserved, 12% of primate-conserved and 10% of all human TAD boundaries to have at least one CTCF binding site overlapping MIR elements (**Fig. 2f-g**). In contrast to ultraconserved boundaries, CTCF sites within the human-specific TAD boundaries were enriched in LTR elements, namely the ERV-K family (**Fig. 2f-g**), which is the most recently endogenized and transcriptionally active ERV family in the human genome^30^. However, these ERVs do not appear to be human-specific, showing that establishment of the human-specific TAD boundaries at these loci has likely followed the TE insertion even. In the mouse genome, the SINE repeat class was significantly enriched (binomial test; p<0.005) at CTCF binding sites of ultraconserved, rodent-conserved and mouse-specific boundaries. Similar to the human genome, among SINEs, the MIR family was significantly over-represented in CTCF binding sites of ultraconserved TAD boundaries, suggesting that this TE family has been involved in the evolution of TAD organization at least since the common ancestor of primates and rodents. Of note, in the mouse genome, the rodent-specific B1 and B2 repeat families were enriched at CTCF binding sites across all evolutionary boundary groups (**Fig. 2f**), supporting the previously proposed hypothesis that many CTCF sites in the mouse genome are derived from B elements^24^. Altogether, our data highlights the correlation between the evolutionary age of TAD boundaries and TEs, suggesting that waves of TE insertions followed by co-option have contributed to the spatial organization of the genome at different evolutionary timepoints.

### Species-specific boundaries are over-represented at evolutionary breaks of synteny

We and others^12,14,31^ have shown that breakpoints of evolutionary chromosomal rearrangements often co-localize with TAD boundaries, which can be interpreted as evidence that selection pressures maintain TAD integrity even when genome synteny is disrupted. We took advantage of our higher resolution Hi-C data and improved genome assemblies^21,32^ (**Supplementary Notes**) to re-assess the extent of the co-occurrence between breakages of synteny (BOS) regions and TAD boundaries across species (**Methods** and **Extended Data Fig. 4**). Consistent with previous reports, the highest number of evolutionary rearrangements was found in gibbon genomes^13,32,33^ (168 BOS in Hylobates and 130 in Nomascus) and the lowest number of BOS was found in Caroli (7 BOS relative to mouse; **Supplementary Table 5**). We observed a strong reduction in genomic interaction frequency across BOS, as well as a notable dip in the collective insulation score at BOS in all three non-human primate species (**Fig. 3a**). Caroli and Pahari showed similar patterns, but the signal was localized and much weaker, likely due the relatively small number of BOS in these species. Consistently, the overlap between BOS and TAD boundaries was statistically significant in all three primates, but not in the rodent genomes (Fisher’s exact test p<0.05; **Fig. 3b** and **Supplementary Table 6**).

**Fig 3.**
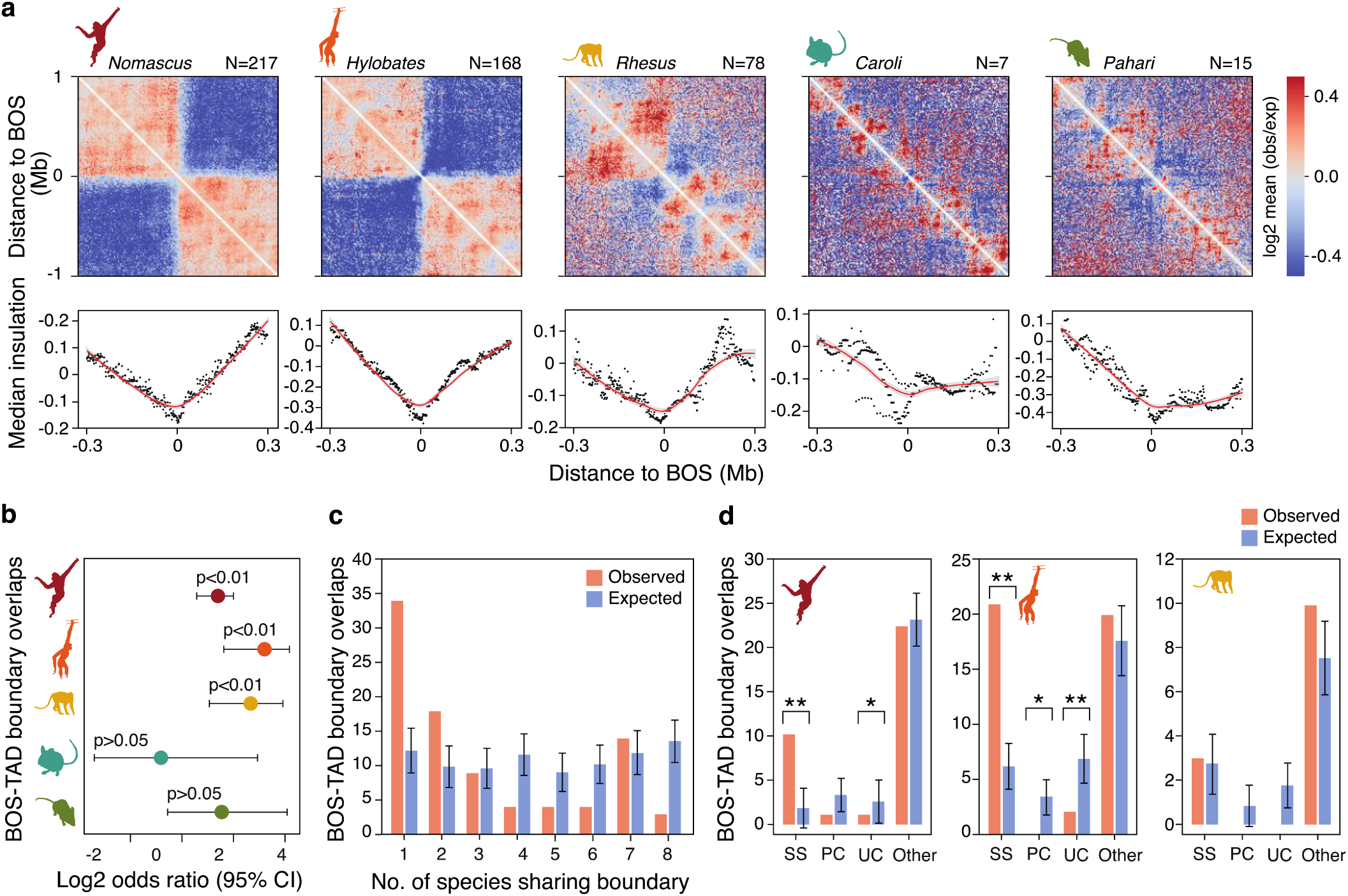
BOS are associated with species-specific TAD boundaries. **a**, Heatmaps show frequency of genomic interactions and dot plots show median insulation scores around BOS. **b**, Log2 odd ratio for the overlap between BOS and TAD boundaries is shown across genomes. **c**, Bars show observed and expected frequency distribution of cross-species conservation levels of all TAD boundary overlapping BOS in rhesus, Nomascus or Hylobates gibbons. **d**, Observed and expected frequency distribution of evolutionary conservation level of TAD boundaries overlapping BOS is shown. **: p<0.005, * p<0.05.

The formation of TAD boundaries that overlap BOS regions could either predate or follow the evolutionary rearrangement event, but thus far, the relative timing of these events has not been investigated. Boundaries predating the rearrangement event are expected to have higher cross-species conservation compared to those established specifically in association to the breakage. We leveraged the large number of BOS and the significant overlap between TAD boundaries and BOS present in the genome of all non-human primates included in our study to investigate these scenarios. A collective investigation of all TAD boundaries that overlap BOS across the three primate species revealed over-representation of species-specific and low-conservation TAD boundaries at BOS (**Fig. 3c**). Consistently, within each of the three species, we observed that ultraconserved boundaries and primate-conserved boundaries were significantly under-represented at BOS, while overlaps with species-specific boundaries were significantly more prevalent than expected in both Nomascus and Hylobates (one-tailed permutation p<0.05) (**Fig. 3d**). In the rhesus genome, TAD boundaries overlapping BOS displayed an overall pattern similar to gibbons, but statistical significance was not reached likely due to the lower number of TAD boundaries and BOS overlaps in this species. Overall, these findings suggest that formation of most TAD boundaries that overlap BOS co-occurred or followed the evolutionary chromosomal rearrangement event.

### Deletion of ultraconserved TAD boundaries results in selective phenotypes

TAD boundaries help prevent ectopic gene regulatory interactions across neighboring TADs, and their disruption can lead to gene mis-regulation of nearby genes^5,15^. As such, there may be an association between TAD boundary conservation level and the nature and function of nearby genes. Specifically, we expect ultraconserved TAD boundaries to be associated with genes whose disruption interferes with fundamental biological functions. Consistently, using the Genomic Regions Enrichment of Annotations Tool (GREAT^34^) to investigate ontology terms associated with TAD boundaries (**Methods**) revealed significant enrichment of genes implicated in ‘anatomical structure formation during embryogenesis’ nearby ultraconserved boundaries in the human genome (GO: 0048646, q<0.05; **Supplementary Table 7**).

To test if ultraconserved TAD boundaries are essential for proper gene regulation and function, we used CRISPR-Cas9 to investigate functionality of two highly conserved TAD boundaries in mice (**Fig. 1a**). These boundaries were selected based on the following criteria: 1) not overlapping protein-coding genes; 2) containing cross-species conserved CTCF binding site; and 3) presence of development-associated genes flanking the TAD boundary. The first deletion targeted a 23Kb region (mm10, chr4:107716841-107739906) between the *Dmrtb1* (known as *Dmrt6* in mouse) and *Lrp8* and included a conserved CTCF binding site ~14Kb downstream of an ultraconserved TAD boundary “B396” (**Fig. 4a**). This deletion strategy was adopted in order to include a BOS between human and Nomascus, since TAD boundaries associated with evolutionary rearrangements may have evolved under additional evolutionary constraints^14^. The *Dmrtb1* gene belongs to the highly conserved DMRT protein family and has been associated with spermatogenic impairment in humans^35^. In mice, this gene plays an essential regulatory role in B spermatogonia in the transition from mitosis to meiosis and activation of meiotic genes^36^. *Lrp8* encodes a low-density lipoprotein receptor highly expressed in testis^37^. We thus focused our analyses on testis-related phenotypes. The deletion was confirmed in mouse lines (*B396_BOS*) via PCR, sequencing, and Southern blots (**Supplementary Notes** and **Supplementary Fig. 2**). Capture Hi-C^38^ on bulk testis did not detect any significant differences in chromatin interaction patterns around the deletion site between homozygous mutant (*B396_BOS^-/-^*) and wild-type mice (*B396_BOS^+/+^*) (**Methods** and **Extended Data Fig. 5a**). However, RNA-seq analysis revealed a significant (p=0.023; Wald test) dose-dependent increase in *Dmrtb1* expression in *B396_BOS^-/-^* compared to *B396_BOS^+/+^* mice, and an increase —albeit not significant (p=0.278; Wald test)— in *Lrp8* expression in heterozygous (*B396_BOS^+/-^*) mice (**Fig. 4b**). To validate these findings, as well as determine if gene expression differences were specific to a given stage of spermatogenesis, we performed RNA *in situ* hybridization with RNAScope using probes against *Dmrtb1* and *Lrp8* in homozygous mutant and wild-type testis. Both transcripts were expressed in different cell types throughout the cycle of the seminiferous tubules grouped into Early (I-V), Mid (VI-early VIII) and Late (late VIII-XII) spermatogenesis stages. Image analysis of testis cross-sections (n=120 tubules) confirmed a small, but significant, increase in expression of *Dmrtb1* (p< 10^-4^; Wilcoxon test) and *Lrp8* (p< 10^-31^; Wilcoxon test; **Fig. 4c-d**).

**Fig 4.**
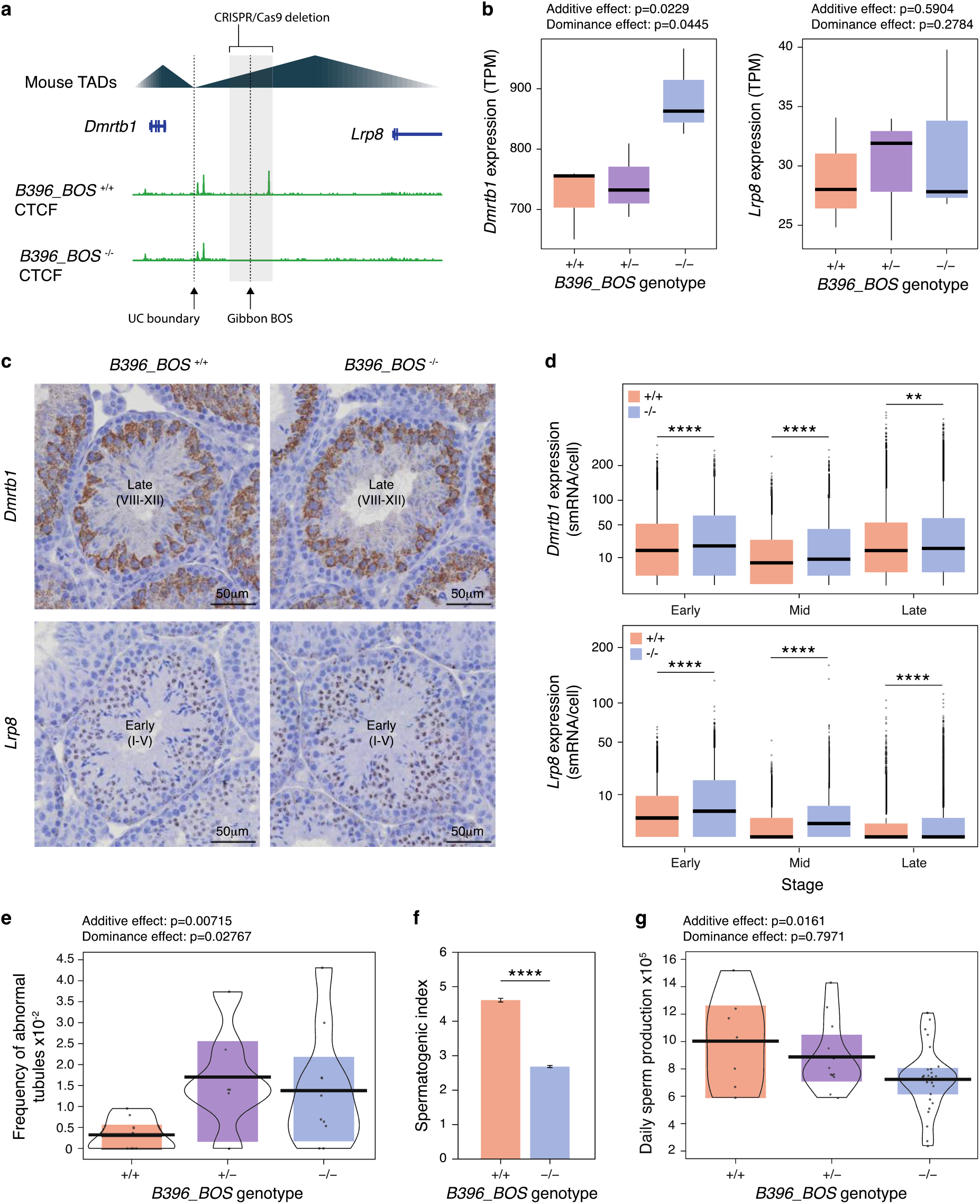
Testis of *B396_BOS^-/-^* mice display changes in testicular *Dmrtb1* expression, histology and function. **a,** A schematic view of the deleted region in B396_BOS-/- mice. **b,** Testicular Dmrtb1 and Lrp8 expression is shown for different genotypes of the B396_BOS line based on RNA-seq data. **c,** *In situ* hybridization (ISH) assay for Dmrtb1 and Lrp8 transcripts in testis of B396_BOS -/- and control mice show representative tubules at identical early (I-V) or late (VIII-XII) stages of spermatogenesis. **d,** Quantification of Dmrtb1 and Lrp8 (n=60 tubules/transcript) signal from the testis ISH assay. **e,** Frequency of abnormalities in testis tubules from different genotypes (n=5,248). **f,** Spermatogenic index is shown as an estimate of testicular function (n= 3,464 tubules). **g,** Daily sperm production estimates are shown (n=38). Abbreviations are as follows: TPM: transcript per million; smRNA: single-molecule RNA; ****: p ≤ 0.0001; **: p ≤ 0.01.

We next investigated whether this deletion caused changes in testis function (**Supplementary Notes**). Manual evaluation of Periodic Acid-Schiff (PAS) stained tubules (n=5,248) revealed an increase in the frequency of abnormal tubules in heterozygous and homozygous mutants compared to wild-type (p= 0.007; Wald test), with evidence for dominance of the mutated allele (p= 0.028; Wald test) (**Fig. 4e**). Furthermore, automated image analysis of testicular cross-sections (n= 3,464 tubules) using SATINN^39^ (**Fig. 4f**) showed a 35% decrease in efficiency of spermatogenesis in homozygous mice (4.05 ± 0.048 to 2.63 ± 0.038, mean ± SE; p< 10^-5^), matching the 28% decrease estimated by daily sperm production (**Fig. 4g**; p=0.016; Wald test) from frozen testis of the same mice. Taken together, these data show an increase in *Dmrtlb* and*Lrp8* expression in *B396_BOS* mutant testis, with subtle quantitative effects on sperm production.

A second deletion of an ultraconserved boundary targeted a ~18Kb region (mm10, chr2:115,840,641-115,858,055) upstream of *Meis2*, a TALE transcription factor involved in inner-ear^40^ and heart development^41–43^ (**Fig. 5a**). We obtained four stable mouse lines (*B5234*) that were validated via PCR, sequencing and Southern blot, and used two of them for downstream phenotyping (**Supplementary Fig. 2**). Due to the known role of *Meis2* in heart development^41–43^, we investigated the consequences of this deletion in this organ first. CTCF ChIP-seq confirmed successful removal of the targeted CTCF binding sites (**Fig. 5a**) and qPCR showed significant upregulation of *Meis2* expression in the heart tissue of *B5234^-/-^* mice (**Fig. 5b**). Visual comparison of cardiac histology (H&E staining) of *B5234^+/+^* and *B5234^-/-^* mice (n=4/group) showed no apparent overt cardiac defects, including no differences in cardiac wall thickness between the left and right ventricles (LV and RV, respectively). However, LV walls in *B5234^-/-^* mice appeared more compact, with less extracellular tissue/space in between cardiac cells, and displayed irregular trabeculation in some regions (**Fig. 5c**). Comparison of extracellular space within the LV cardiac wall revealed that *B5234^+/+^* mice exhibited a significantly larger percentage of extracellular space than *B5234^-/-^* mice (Student’s t-test; p = 0.00004) (**Fig. 5d**).

**Fig 5.**
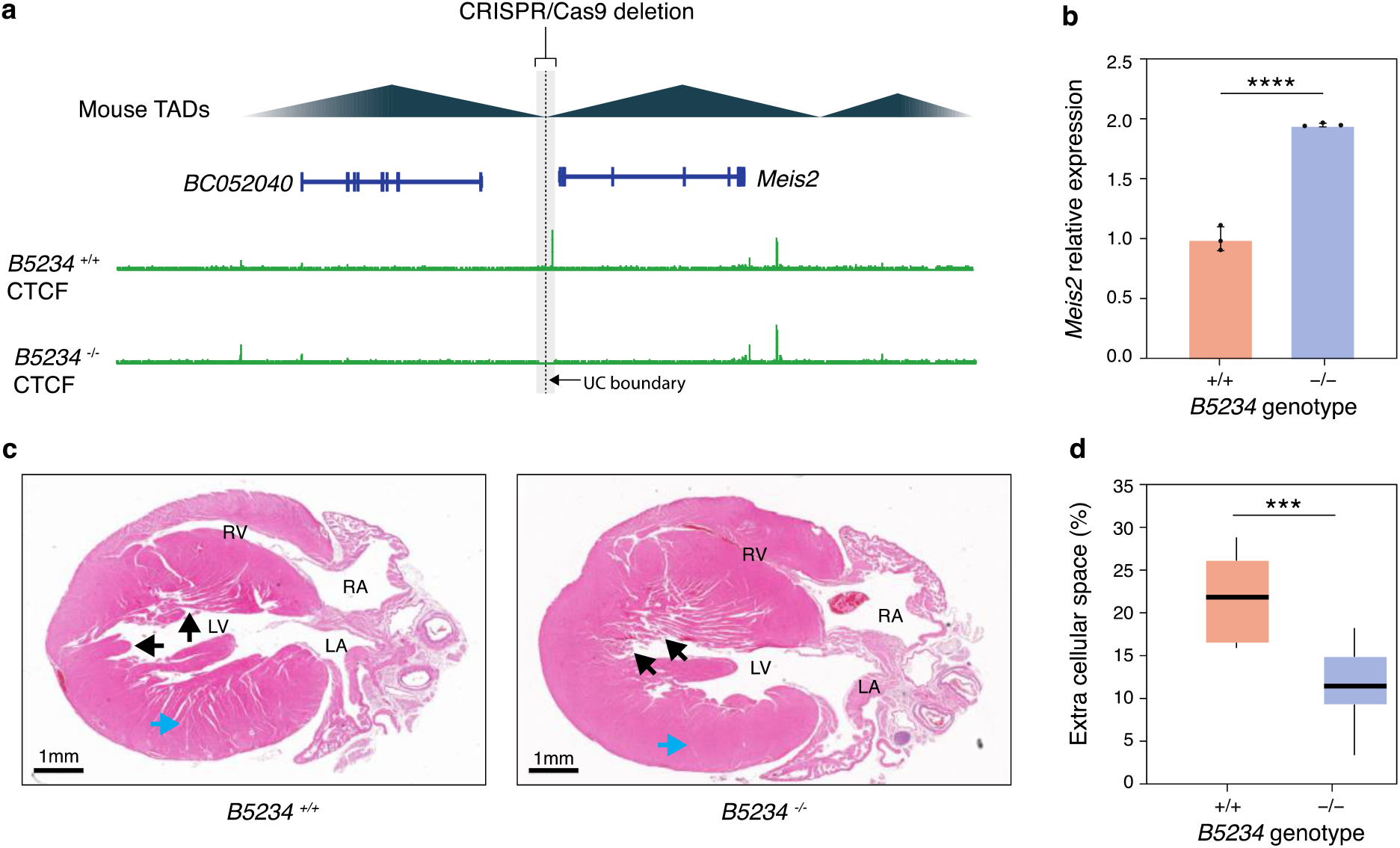
Hearts of *B5234^-/-^* mice exhibit changes in *Meis2* gene expression and histology. **a**, Schema of the deletion in *B5234^-/-^* mice. **b**, RT-qPCR reveal increased *Meis2* expression in hearts of *B5234^-/-^* mice. **c**, H&E staining of longitudinal sections of *B5234^-/-^* and *B5234^+/+^* mouse hearts show differences in LV compaction (blue arrow) and trabecular structures (black arrows). **d**, Quantification of extra-cellular space from H&E images confirms increase compaction (decrease extra-cellular space) in hearts of *B5234^-/-^* in comparison to *B5234^+/+^* mouse hearts. (Abbreviations are as follows: LV: Left ventricle; RV: Right ventricle; LA: Left atrium; RA: Right atrium; ****: p<0.00001, ***: p<0.0001).

Given *Meis2’s* role in ear development^40^, we also characterized the morphology of the utricle and the cochlea in neonatal mice (**Supplementary Methods**) and found that these structures remained unchanged among all three genotypes (*B5234^+/+^, B5234^+/-^, and B5234^-/-^*). Hair cells appeared to have normal morphology and bundle orientation in both the cochlea and utricle (**Supplementary Fig. 3**). Consistent with these findings, no differences were found in the hearing levels between the three genotypes measured by auditory brainstem response (ABR) recordings in seven-month-old mice (**Supplementary Methods**). Overall, deletion of the two ultraconserved boundaries resulted in gene expression changes and mild, but significant, phenotypic changes relevant to the biological functions of the genes nearby the boundary.

### Deletion of a human-specific boundary alters gene expression in human neurons

Emergence of new TAD boundaries changes local TAD organization, which could lead to the evolution of novel and adaptive gene regulatory modules. To this end, among the different boundary conservation groups, the human-specific boundaries are of particular significance as they may be relevant to evolution of human-specific traits and adaptations. When investigating gene ontologies (GO) associated with human-specific TAD boundaries, we detected a significant enrichment of pathways pertaining to ‘positive regulation of synapse assembly’ (GO:0051965, q<0.05; **Supplementary Table 8**). Synapse formation is one of the main brain development processes that sets humans apart from other primates^44^ and its pathology sits at the heart of several human cognitive disorders, such as autism spectrum disorders (ASD) and schizophrenia^45^. Human-specific boundaries were also associated with several genes implicated in brain disease and development, such as the Autism Susceptibility Candidate 2 (*AUTS2*) gene, a key regulator of transcriptional networks and a mediator of epigenetic regulation in neurodevelopment. *AUTS2* is implicated in ASD and other neurological diseases^46, 47^ and contains the most significantly accelerated genomic region differentiating humans from other primates^46^. To further investigate the association of human-specific TAD boundary deletions with human disease, we tested their overlap with pathological copy number variants (CNVs), previously identified from 29,085 children with developmental delay and 19,584 healthy controls^48^. Using permutation analysis (**Supplementary Notes**), we found significantly higher recurrency of human-specific TAD boundary deletions in children with developmental delay (n=335 out of 1,130 human-specific boundaries), compared to healthy controls (n=161) (Fisher’s exact test, p < 0.0001).

We next characterized the function of a human-specific boundary (B14804) in human neurons located ~200Kb upstream of *AUTS2*, which has been implicated in numerous human neurological diseases^46, 47^, by using CRISPR-Cas9 to delete an 11Kb region (hg38: chr7:69,393,633-69,405,124). The deletion was performed in induced pluripotent stem cells (iPSC) WTC11 Ngn2 cell line, which allows quick and robust differentiation to neurons^49^ (**Fig. 6a**). We obtained two lines carrying independent deletions (*B14804^-/-^* cell line 1 and 2) and validated them via PCR (**Methods and Supplementary Table 9**). Both lines, along with the wild-type line, as a negative control, were differentiated into dopaminergic neurons, as confirmed by expression of neuronal marker genes *MAP2* and *TUBB3* (**Fig. 6b**). We found *AUTS2* expression to be significantly increased in both lines compared to wild-type cells (p=0.0031; Student’s t-test), while no changes were detected in expression of the *HPRT1* housekeeping gene (**Fig. 6b**). We also carried out Hi-C on the two *B14804^-/-^* and wild-type cell lines but did not detect significantly different Hi-C interaction loops between genotypes. However, we observed notably lower insulation scores at the *AUTS2* locus in *B14804^-/-^* cells, compared to wild-type (**Fig. 6c**). Altogether, our findings not only hints at the potential involvement of human-specific TAD reorganization in the evolution of the complex human brain, but also suggests that disruption of human-specific TAD boundaries could contribute to developmental and neurological disorders.

**Fig 6.**
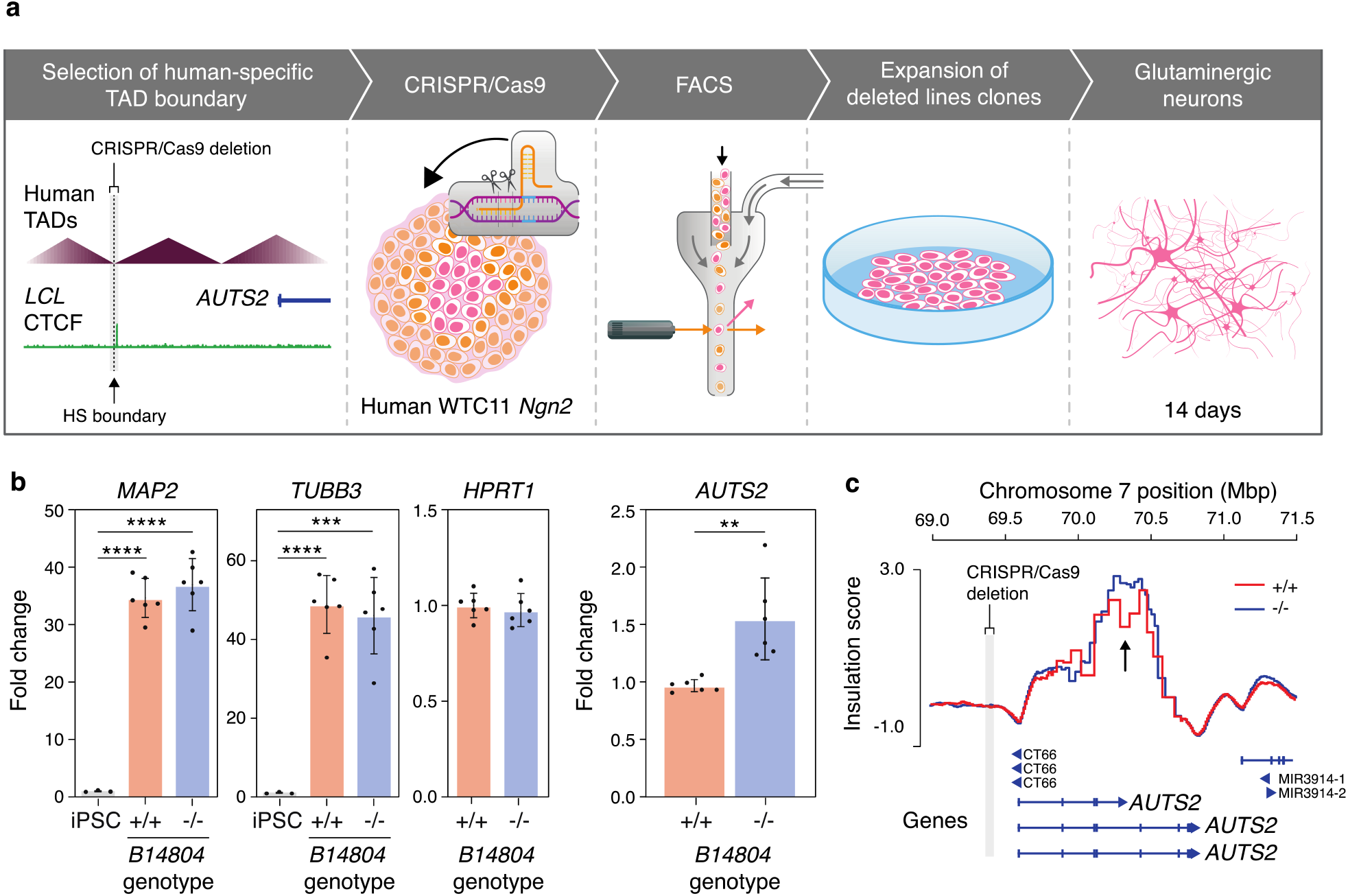
iPSC-differentiated *B14804^-/-^* neurons have lower *AUTS2* expression and a higher local insulation score. **a**, Flowchart shows steps involved in generating *B14804^-/-^* neurons. **b**, Differentiated *B14804* cells show higher expression of neuronal markers, *MAP2* and *TUBB3. B14804^-/-^* neurons show no change in *HPRT1* expression but have higher *AUTS2* expression. **c**, local increase in insulation score is observed at the *AUTS2* locus, downstream of the *B14804* deletion.

## DISCUSSION

In this study, we investigated mammalian TAD conservation by obtaining and comparing high-resolution Hi-C and ChIP-seq data across four primate and four rodent species (**Fig. 1a**). We found that only 14% of human and 15% of mouse TAD boundaries were conserved across all species, which is lower than the conservation level reported between human and mouse^7^. Of note, our cross-species Hi-C datasets originated from two different cell types, LCL and liver, and thus any boundary identified as ultraconserved had to be developmentally stable between these two tissues, adding constraints to our classifications criteria. Nevertheless, our findings challenge the assumption that TAD organization is mostly conserved across tissues and species and show that only a relatively small portion of TAD boundaries present in the genome of the common ancestor of the Euarchontoglires (or Supraprimates) has evolved under strong evolutionary constraints.

Besides establishing the level of TAD conservation, we also explored whether genetic and epigenetic features of TAD boundaries vary across evolutionary groups. We observed that ultraconserved TAD boundaries show higher insulation strength and more CTCF binding sites than species-specific boundaries. These findings are in line with a previous study that reported higher CTCF binding site clustering and stronger CTCF affinity at TAD boundaries shared across five murine species, compared to boundaries with lower conservation^50^. Moreover, CTCF binding sites at ultraconserved boundaries overlap with older transposable elements (namely MIRs), whereas human-specific boundaries are associated with more recent families (ERVs), reinforcing the notion that different waves of transposable elements have been shaping genome topology through insertion of CTCF binding sites at different evolutionary times^24,26,29^. Altogether, this shows that recurrent emergence and maintenance of several, possibly redundant, CTCF binding sites contribute to the higher conservation and insulation strength of ultraconserved TAD boundaries.

TAD boundaries play an important role in regulating expression of nearby genes^51^, and disruption of boundaries can lead to gene mis-regulation and aberrant phenotypes^4–6,52^. Hence, it is likely that TAD boundaries evolve under strong evolutionary constraints preventing their disruption, even in face of evolutionary chromosomal rearrangements. In line with the expectation that TAD boundaries evolve under strong evolutionary constraints, we would expect structural variations that disrupt TADs to be under purifying selection. As a result, evolutionary breaks of synteny (BOS) should co-localize with TAD boundaries to keep TADs intact^12,14,53,54^. Here, we report significant co-localization between BOS and TAD boundaries in the gibbon species, which carry highly rearranged genomes^13^, and a similar —albeit weaker— BOS-boundary correspondence in the rhesus macaque. Additionally, our evolutionary classification enabled us to detect an over-representation of species-specific (i.e., more recent) TAD boundaries among those that co-localize with BOS **(Fig. 3c-d)**, suggesting that evolutionary chromosomal rearrangements likely co-occur within novel TAD structures and regulatory modules, rather than predating them. Hence, these findings suggest a connection between the timing of genome reshuffling and TAD boundary establishment.

Another implication of the functional connection between TAD boundaries and genes, is that selection pressures under which TAD boundaries evolve are likely influenced by the function of nearby genes. Consistently, we observed that genes associated with ultraconserved boundaries in the human genome were enriched in pathways pertaining to anatomical structure formation during embryogenesis, whose disruption is likely to lead to developmental defects. Moreover, manipulating two ultraconserved TAD boundaries in mouse models altered expression of neighboring genes and resulted in phenotypic consequences that could lower reproductive and/or survival fitness. While our deletions resulted in relatively mild phenotypes, a recent study^19^ performing large (35.3 + 26.5Kb; mean + stdev) *in vivo* deletions of TAD boundaries near developmental genes in mice observed severe phenotypes when two boundaries classified as ultraconserved in our dataset were deleted. Of note, the severity of phenotypes reported in this study seems to broadly correlate with the deletion size and number of CTCF clusters removed. As our deletions were smaller in size and only removed one or two CTCF binding sites, it is possible that retention of functionally redundant CTCF binding sites mitigated effects of the deletions at ultaconserved boundaries.

We also identified TAD boundaries exclusively present in the human genome and showed that they were significantly associated with genes implicated in synapse assembly. The human brain undergoes unique synapse patterning, which is thought to contribute to its extraordinary complexity and connectivity^44^. For instance, timing of synaptic development and expression of relevant genes is prolonged in humans compared to macaque and chimpanzee, and synapse density is highly increased in humans in comparison to other primates^55^. Importantly, mounting evidence also places synapse assembly and its pathology at the heart of several human cognitive disorders, such as ASD and schizophrenia^45^, suggesting that disruption of human-specific boundaries might also be a contributing factor in human neuropathology. It should be noted however, that this study used Hi-C data from liver and LCL tissues to identify human-specific boundaries, and given that TAD organization can vary across tissues^56^, association of human-specific boundaries with brain-related genes and pathways will likely be stronger if investigated directly in brain cells^56^. Nevertheless, we found significantly more overlap between human-specific TAD boundaries and CNVs identified in children with developmental delay^48^, compared to healthy controls. This is in line with recent work that found TAD-shuffling to alter enhancer-promoter interactions and be associated with a variety of human developmental disorders^57^. We additionally showed that deletion of a candidate human-specific TAD boundary upstream of the *AUTS2* gene resulted in notable differences in genomic interaction patterns (i.e., insulation strength) and gene expression at the *AUTS2* locus in iPSC-derived human neurons, providing evidence for functionality of human-specific boundaries in human brain cells.

In summary, we generated high-resolution Hi-C and ChIP-seq data from several species and used direct multi-species comparison to stratify TAD boundaries in the mouse and human genomes based on evolutionary conservation. Our study highlights the remarkable range of diversity that exists in the evolutionary conservation of TAD boundaries, regardless of species (mouse vs. human) and tissue (LCL vs. liver). Furthermore, by comparing genetic and epigenetic features of boundaries across conservation groups and investigating function of candidate TAD boundaries via *in vivo* and *in vitro* manipulations we provided new insights into the functional implications of TAD boundary evolution. While our study underlines the importance of TAD conservation in maintaining proper gene regulation patterns during evolution, it also showcases how recent changes in TAD organization can contribute to emergence of evolutionary novelties and species-specific traits. Our findings also showcase the importance of our evolutionary TAD boundary annotations in interpreting genetic data from clinical cases and should be utilized to assess potential disease-association with TAD boundary altering CNVs. Future comprehensive comparative studies in additional tissues and developmental stages will help us help us better understand contributions of TAD organization to biological function, as well as if and how disrupting these structures can lead to pathology.

## METHODS

### Hi-C library generation, TAD identification and evolutionary classification

We used liver tissue and lymphoblastoid cell lines (LCLs) to generate Hi-C libraries with the Arima Hi-C Kit following manufacturer’s protocol. Briefly, ~5ug frozen pellets of the homogenized fixed liver or LCL were lysed and conditioned before chromatin digestion. The digested chromatin was then filled in and biotinylated before ligation. Next, chromatin was protein-digested and reverse crosslinking overnight, followed by purification. The purified DNA was then sonicated using the bioruptor pico (Diagenode) and size selected before library preparation using the NEB DNA Ultra II, following Arima’s protocol. The Hi-C DNA was bound to streptavidin beads before enzymatic end prep, adaptor ligation, DNA release by heat incubation, and lastly, PCR to barcode and amplify the libraries. Libraries were sequenced on the Illumina HiSeq2500 or NovaSeq6000. Raw Hi-C sequencing was processed using the HiCUP pipeline^58^. Alignments were converted to a multi-resolution Hi-C matrix using pairtools (https://github.com/mirnylab/pairtools). Hi-C data between biological replicates were combined after verifying their correlation based on the Pearson correlation as calculated by the HiCExplorer package^59^ (**Extended Data Fig. 1**). TAD boundaries were called at 10Kb resolution using the hicFindTADs command from HiCExplorer using the flags “--minDepth 100000 and –maxDepth 600000” (**Supplementary Table 1**). Briefly, this program uses running windows of different sizes to measure interaction separation or “insulation score”) between the two sides of each Hi-C matrix bin. The “insulation score” describes the degree of interaction separation across the Hi-C matrix by calculating the mean z-score of contacts across a genomic locus. Smaller insulation score values indicate stronger insulation at that position in the Hi-C matrix. TAD boundaries are then determined as bins with a statistically significant local minimum insulation score.

In order to perform cross-species comparisons and determine TAD boundary evolutionary conservation, we lifted TAD boundaries identified in all species to the hg38 coordinates, generated a union boundary map and used paradigms of boundary presence/absence across species to stratify union boundaries into groups, as detailed in the **Supplementary Notes**.

### Epigenetic and genetic characterization of TAD boundaries across conservation groups

We compared insulation score, chromatin structure, CTCF binding sites, and overlap with genes and transposable elements (TE) across TAD boundary conservation groups in human and mouse genomes. Boundary insulation scores were generated as part of the TAD boundary annotation analysis and compared using the Wilcoxon signed-rank test. CTCF binding sites and chromatin states were determined by using a combination of public and newly generated H3K4me1, H3K27ac, K3K4me3, H3K27me3 and CTCF ChIP-seq data from a female and a male of each of the eight species examined in this study (**Supplementary Table 2**). All rodent and rhesus ChIP-seq libraries were generated from liver tissue, while lymphoblastoid cell lines (LCL) were used for the rest of the species. ChIP-seq libraries were generated and analyzed as described in the **Supplementary Notes**. Refseq hg38 and mm10 gene annotations were intersected with TAD boundary coordinates, using bedtools^60^, to determine proportion of boundaries overlapping with genes. We used the TEanalysis tool^61^ to examine TE enrichment at CTCF ChIP-seq peaks (i.e. binding sites) within TAD boundaries in each conservation group.

### Investigating overlap of breaks of synteny (BOS) with TAD boundary groups

We identified all breaks of synteny (BOS) of evolutionary chromosomal rearrangements using a custom pipeline in which all pairwise comparisons of known chromosomes of the “target” genome are aligned to the “query” genome using LASTZ (https://github.com/carbonelab/lastz-pipeline), followed by alignment chaining and filtering using UCSC tools^61^. For Nomascus, we took advantage of an improved version of the genome assembly (Asia_NLE_v1) based on PacBio CLR and guided by Nleu3 generated by the Eichler lab (**Supplementary Notes**). Pairwise genome alignments were then processed using a custom python script (https://github.com/carbonelab/axtToSyn), which elongates alignment blocks that are longer than 1Kb and have a minimum alignment score of 100,000, by merging them with other blocks on the same chromosome and strand. Elongated alignment blocks represent synteny blocks between two genomes, thus synteny breakpoints were defined as 1Kb regions flanking each elongated synteny block in the target genome. To transfer coordinates of synteny breakpoints from the target genome coordinates to those of the query genome, we used BLAT^62^ with the following parameters: -stepSize=5 - repMatch=2253 -minScore=20 -minIdentity=0. The BLAT results were manually evaluated and if the BLAT score of the second highest-scoring hit was within 10% of the top-scoring hit, the breakpoint was annotated as duplicated and removed. Breakpoints that survived this filtration step were further manually inspected to remove BOS overlapping segmental duplications and large repeats. Only curated breakpoints that survived both filtration steps were used for downstream analysis (**Supplementary Table 5**).

Breaks of synteny (BOS) were identified in each of the five species pairs using a custom R-script as described in the **Supplementary Notes**. Using custom scripts (https://github.com/carbonelab/hicpileup) we visualized aggregate Hi-C contact frequencies in a 2Mb window centered at synteny breakpoints. Median insulation score was visualized in 600 Mb windows centered at synteny breakpoints. To test the statistical significance of the overlap between BOS and TAD boundaries we performed permutation tests by using “bedtools shuffle” to shuffle BOS within their original chromosome and used the Fisher’s exact test to compare the number of synteny breakpoint/TAD boundary overlaps (at least 1bp) in shuffled dataset to the true observations across species (**Supplementary Table 6**). We next identified all TAD boundaries located within 30Kb of a BOS in the genomes of species that showed significant overlap between BOS and TAD boundaries (two gibbons and rhesus). In each species, we matched each overlapping boundary to its corresponding union boundary and used the presence/absence in other species to identify species-specific, primate-conserved and ultraconserved boundaries. For each species we then compared the number of boundaries in each conservation group to expectation, which was determined based on randomly selecting the same number of TAD boundaries. We repeated this process 100 times and in each boundary group, the proportion of times out of 100, when the observed number of TADs were more extreme than random expectation was the empirical one-tailed p-value (**Fig. 4c-d**).

### Generation of knockout mice

All transgenic animal experiments were conducted in accordance with the Guide for the Care and Use of Laboratory Animals established by the National Institutes of Health. Protocols were approved by Institutional Animal Care and Use Committees (IACUC) at Oregon Health and Science University (OHSU) and UCSF. All mice were allowed ad-libitum access to food and water and were maintained on a 12h light/dark cycle in a climate-controlled facility. Our first candidate conserved boundary (union_B396) overlapped a BOS region in the *Nomascus leucogenys* gibbon genome and a nearby conserved CTCF peak. Knockout mice harboring deletions of this boundary (*B396_BOS^-/-^*) were generated at the Transgenics Mouse Models core at OHSU by targeting a 23Kb region (chr4: 107716841-107739906) as described in the **Supplementary Notes**. Our second CRISPR deletion (*B5234^-/-^*) targeted an 18Kb region (chr2: 115,840,641-115,858,055, mm10) including an ultraconserved boundary near the *Meis2* gene using i-GONAD^62^ described in **Supplementary Notes**. Offspring from both assays were screened for the deletion using Southern blot, sequencing and a custom designed PCR assay, with primers flanking the deletion site (**Supplementary Table 9**). Molecular phenotyping and histology assays for the mice are described in detailed in the **Supplementary Notes**.

### Generation and phenotyping of the *B14804^-/-^* knockout cell line

We performed CRISPR knockout assays targeting an 11Kb region (hg38: chr7:69,393,633-69,405,124) that included the human-specific B14804 boundary in WTC11-ngn2 cells^49^ (i.e., WTC11 cells with a doxycycline-inducible mouse Ngn2 transgene). Briefly, WTC11-ngn2 cells were cultured in mTeSR1 media (STEMCELL Technologies) with daily media changes following normal WTC11 maintenance protocols. Cells were seeded at a density of 300k cells per 6-well in mTeSR1 media plus Rock Inhibitor (Selleckchem) and cultured for one day. WTC11-ngn2 p37+21 cells (p37= passage number before ngn2 introduction, +21= passage number after the ngn2 insertion) were then transfected with 800ng of each of the four sgRNAs **(Supplementary Table 9)**, 6250ng of TrueCut Cas9 Protein v2 (Invitrogen), and 500ng of MSCV Puro-SV40:GFP plasmid (Addgene) using Lipofectamine CRISPRMAX Cas9 Transfection Reagent (Thermo Scientific) following the manufacturer’s protocol. On the second day post transfection, cells were washed in 1X PBS, dissociated from the plate using Accutase (STEMCELL Technologies), quenched with 1X PBS, spun down and resuspended in a FACs buffer consisting of 1X PBS, 0.5M EDTA (Neta Scientific), 1M HEPES PH7.0 (Neta Scientific), 1% FBS, and Rock Inhibitor. Cells were filtered through a cell strainer, then GFP positive single cells were sorted on a BD FACSAria Flow Cytometer or equivalent using a 100-micron nozzle into 96-well plates containing mTeSR media supplemented with Rock Inhibitor, 1% Penicillin-Streptomycin (ThermoFisher), and 10% CloneR2 (STEMCELL Technologies). Individual colonies were expanded incrementally when wells became confluent. DNA was extracted from a subset of cells of each colony using AllPrep DNA/RNA Mini kit (Qiagen). To validate the deletions, gDNA was extracted using AllPrep DNA/RNA Mini kit (Qiagen), followed by genotyping of each colony using KOD One PCR Master Mix (DiagnoCine) with two unique primer sets (**Supplementary Table 9**). Passage 37+27 WTC11-ngn2 cell lines were differentiated into day 14 neurons following a previously described protocol. In short, cells were seeded and grown in pre-differentiation media. On the third day, cells were dissociated, counted, and plated in differentiation media according to recommended seeding densities. Cells were grown for 14 days, with only a partial media change on day 7. On day 14, DNA and RNA were extracted from the *B14804^-/-^* and wild-type cells using AllPrep DNA/RNA Mini kit (Qiagen). cDNA was synthesized from extracted RNA using SuperScript™ III Reverse Transcriptase (Invitrogen) following manufacturer’s protocol. cDNA was diluted 1:10 and used for qPCR with SsoFast EvaGreen Supermix (BioRad). qPCR reactions were done in triplicate and normalized against *Gapdh* (**Fig. 6b**).

## DATA ACCESSIBILITY

All Hi-C, ChIP-seq, RNA-seq and Capture Hi-C data generated as part of this project is available at the Gene Expression Omnibus (GEO GSE197926). The Nomascus assembly Asia_NLE_v1 is available on NCBI: https://www.ncbi.nlm.nih.gov/data-hub/genome/GCF_006542625.1/

## ACKNOWLEDGMENTS

The ChIP-seq assays were performed by the KCVI Epigenetics Consortium at OHSU. Next-generation libraries were sequenced at the OHSU Massively Parallel Sequencing Shared Resource, the Genomics and Cell Characterization Core Facility at University of Oregon and Novogen Co. Data analyses were performed on the Exacloud computer cluster at OHSU. The ONPRC Integrated Pathology Core provided support services for the research. WTC11-ngn2 cells were a kind gift from Li Gan (Gladstone Institute). We thank Drs. Kent Thornburg and Alina Maloyan for reviewing heart images. We additionally thank Drs. Fudenberg and Pollard for sharing the CNV calls from patients and controls. This work was supported in part by National Human Genome Research Institute (NHGRI) Grant R01HG010333 (for L.C and N.A.), and L.C. and D.F.C. are supported by the National Institute of Health Office of Directors (NIH/OD) Grant P51 OD011092 (to the Oregon National Primate Research Center). B.N. is supported by S10OD021717-01A1 Grant (PI: M. Calvert). E.E.E is supported by National Institutes of Health Grant R01HG002385.

E.E.E. is an investigator of the Howard Hughes Medical Institute. Part of this work was supported by the OHSU University Shared Resources shared grant.

