## Supplementary figures and images for "TAD Evolutionary and functional characterization reveals diversity in mammalian TAD boundary properties and function"

### Extended Data Fig 1

##### Extended Data Fig. 1: Characterization of TAD boundaries in eight different species

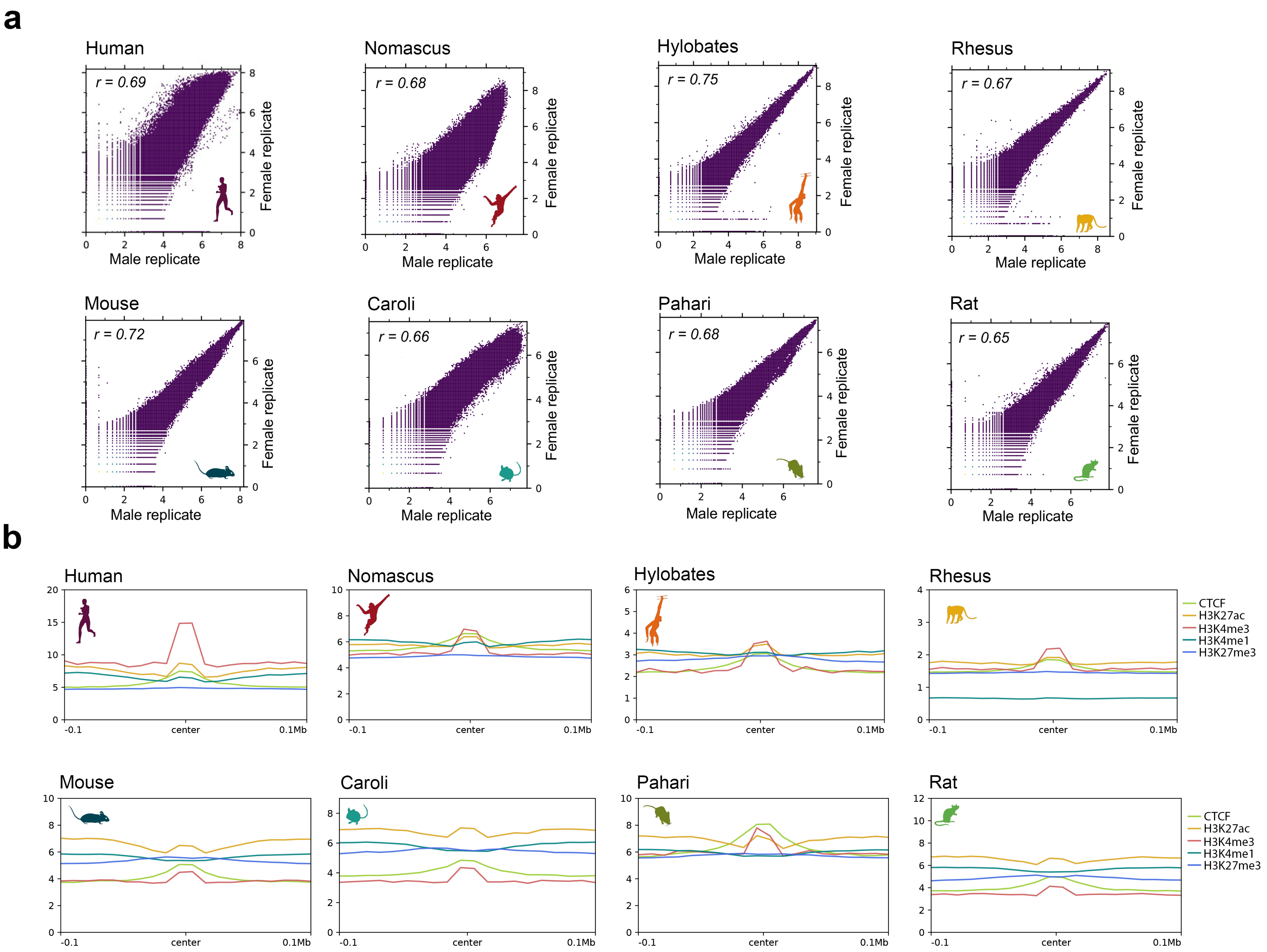

### Extended Data Fig 3

#### Extended Data Fig. 3: Examples of ultraconserved TAD boundaries

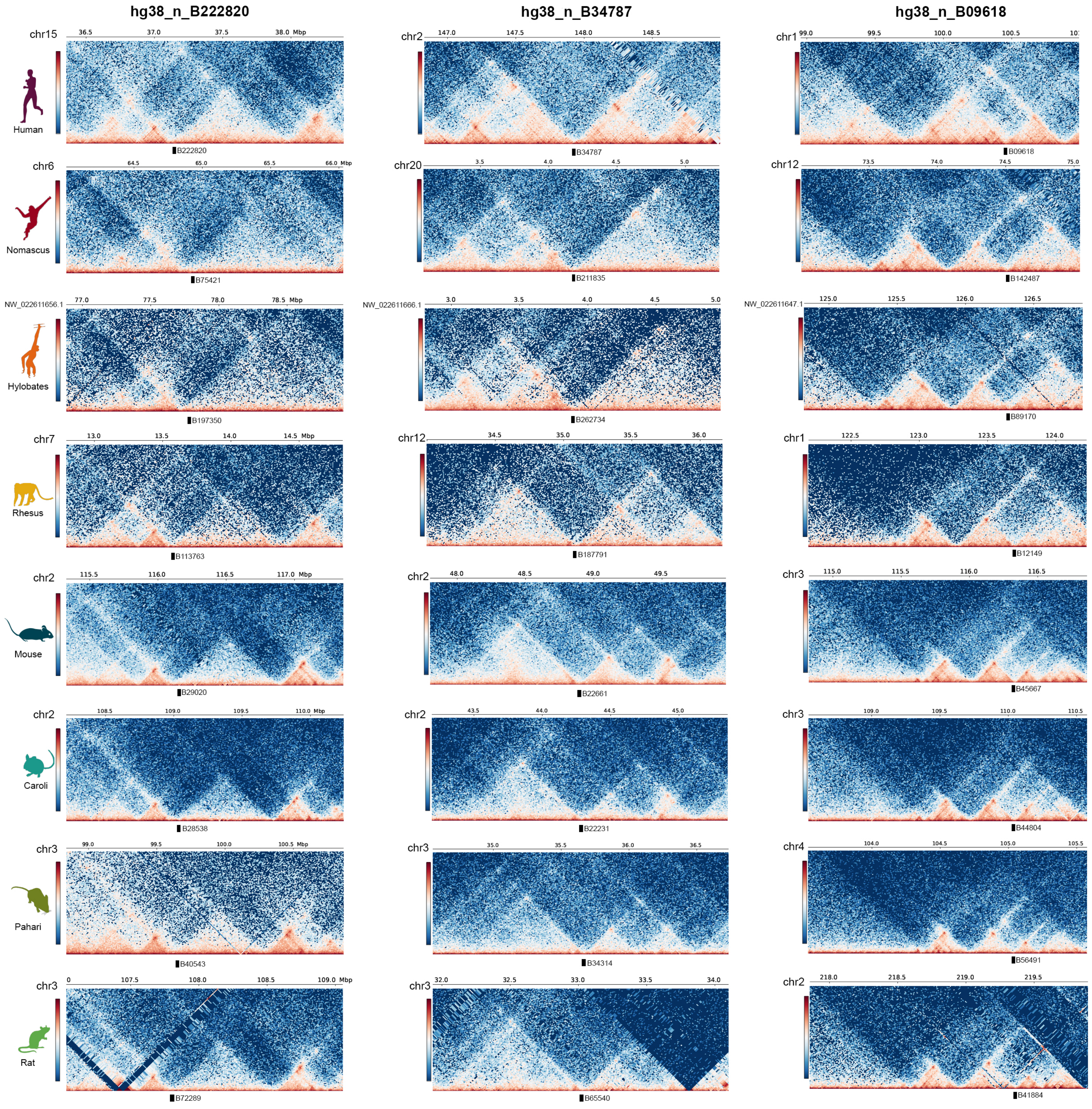

### Extended Data Fig 4

**Extended Data Fig. 4: Pairwise comparisons to identify breaks of synteny (BOS) between species**

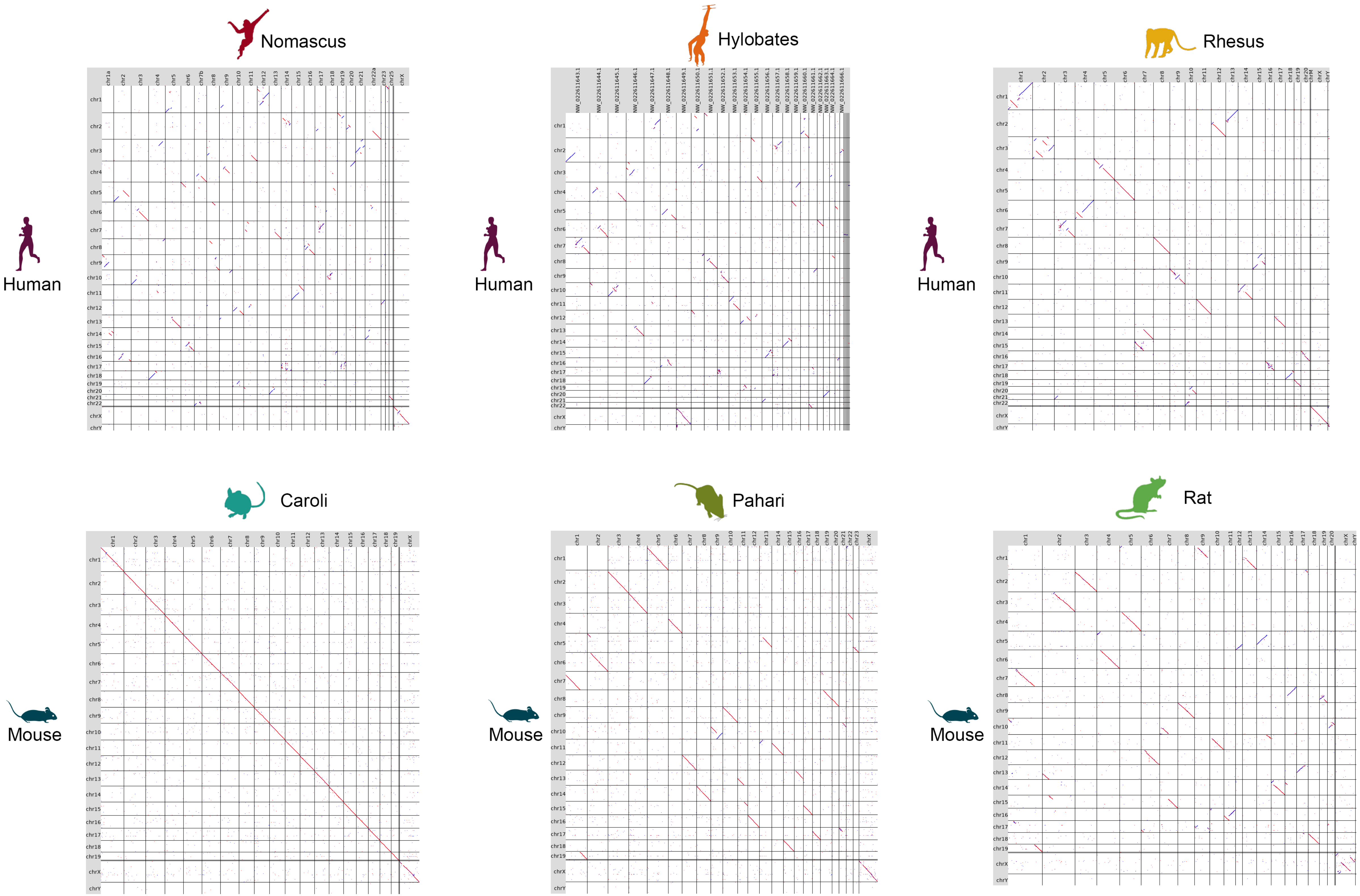

### Extended Data Fig 5

##### Extended Data Fig.5: Phenotyping of B396_BOS-/- mice

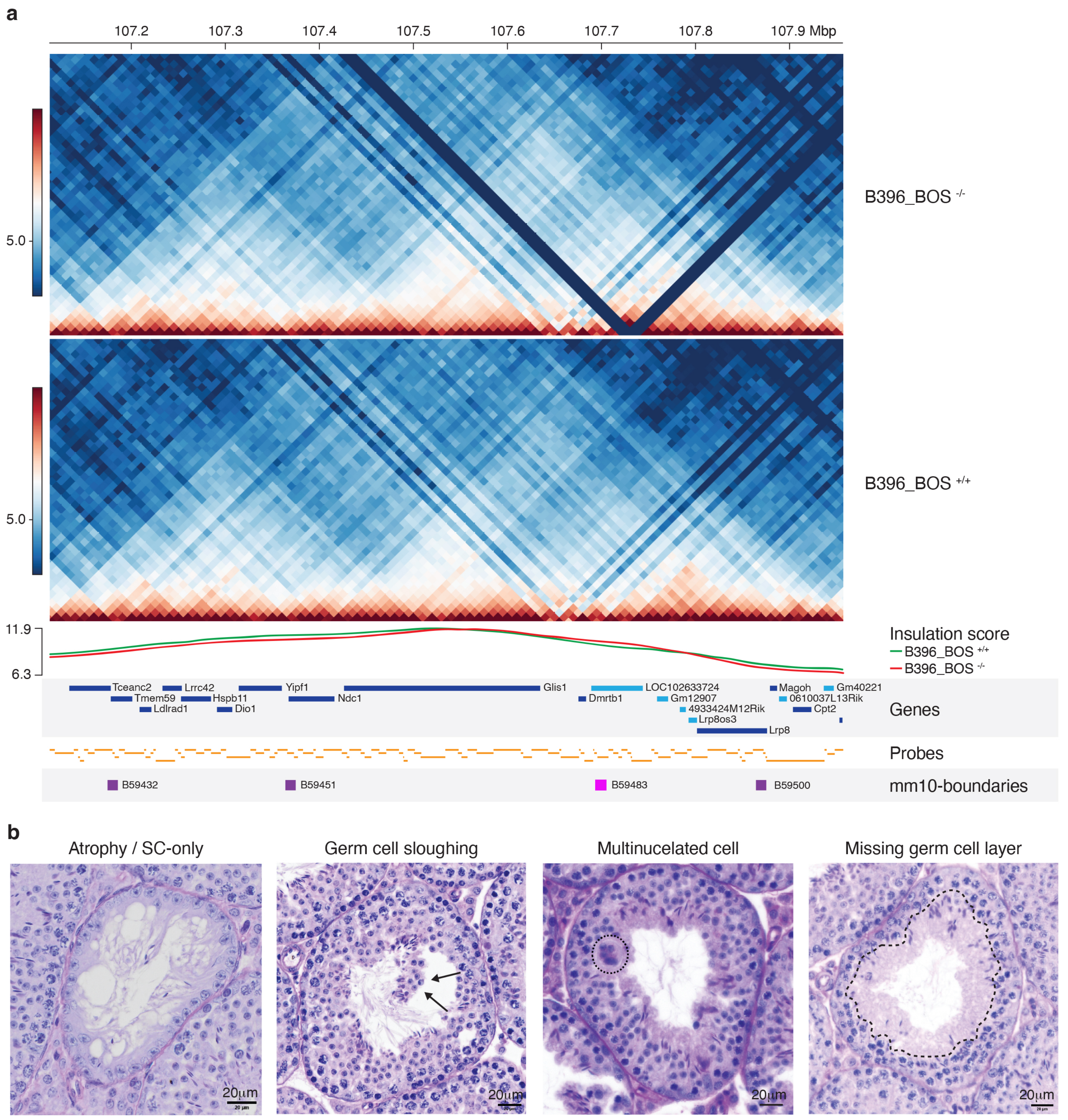
