## Extended Data Fig 2 for "TAD Evolutionary and functional characterization reveals diversity in mammalian TAD boundary properties and function"

**Extended Data Fig. 2: Transposable elements (TE) content at CTCF binding sites in TAD boundaries of each species**

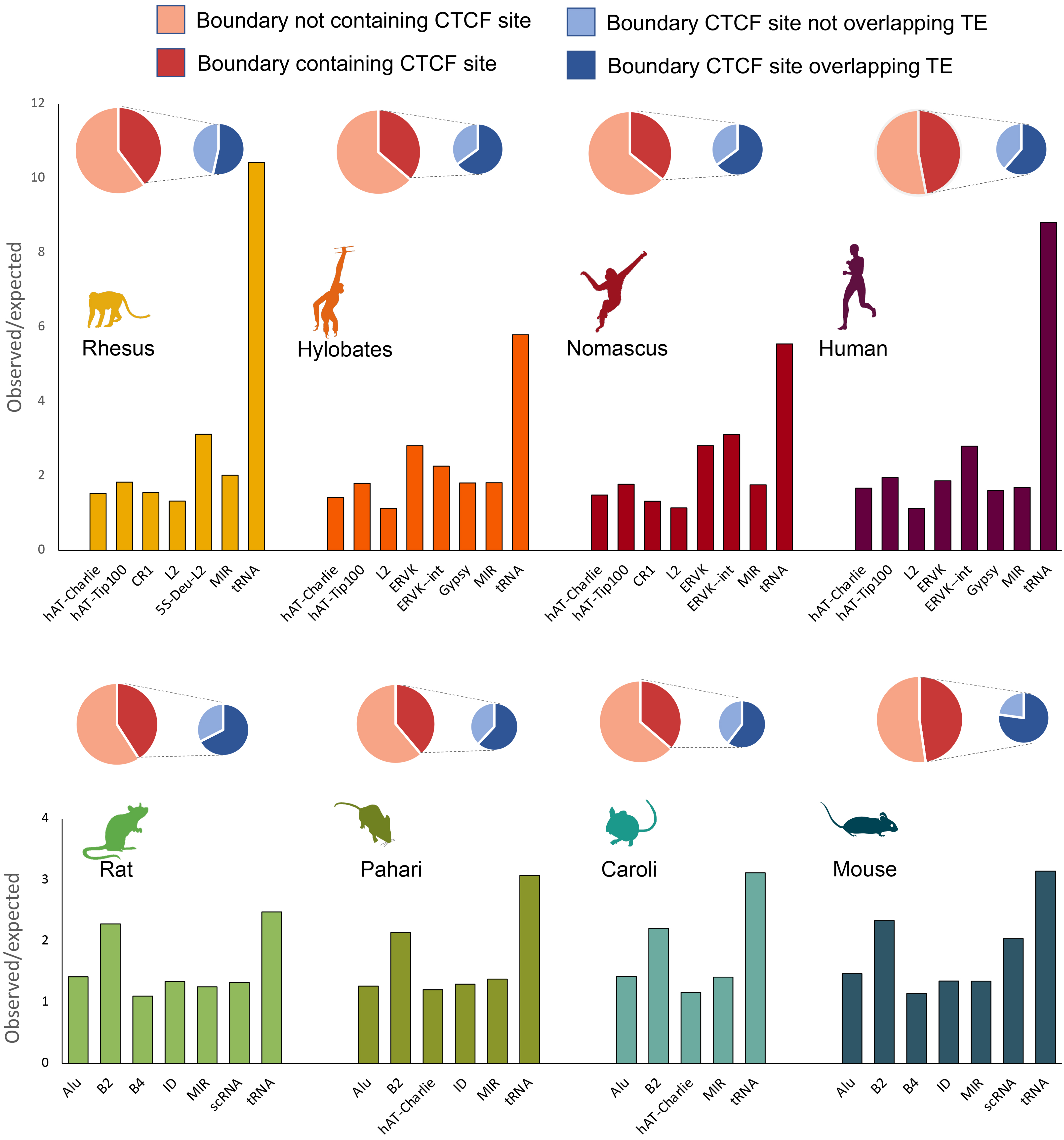
